# A Unified Theory of Response Sparsity and Variability for Energy-Efficient Neural Coding

**DOI:** 10.1101/2024.09.25.614987

**Authors:** Mingyi Huang, Wei Lin, Anna Wang Roe, Yuguo Yu

## Abstract

Understanding how cortical neurons use dynamic firing patterns to represent sensory signals is a central challenge in neuroscience. Decades of research have shown that cortical neuronal activities exhibit high variance, typically quantified by the coefficient of variation (CV), suggesting intrinsic randomness. Conversely, substantial evidence indicates that cortical neurons display high response sparseness, indicative of efficient encoding. The apparent contradiction between these neural coding properties—stochastic yet efficient—has lacked a unified theoretical framework. This study aims to resolve this discrepancy. We conducted a series of analyses to establish a direct relational function between CV and sparseness, proving they are intrinsically correlated or equivalent across different statistical distributions in neural activities. We further derive a function showing that both irregularity and sparsity in neuronal activities are positive functions of energy-efficient coding capacity, quantified by Information-Cost Efficiency (ICE). This suggests that the observed high irregularity and sparsity in cortical activities result from a shared mechanism optimized for maximizing information encoding capacity while minimizing cost. Furthermore, we introduce a CV-maximization algorithm to generate kernel functions replicating the receptive fields of the primary visual cortex. This finding indicates that the neuronal functions in the visual cortex are optimal energy-efficient coding operators for natural images. Hence, this framework unifies the concepts of irregularity and sparsity in neuronal activities by linking them to a common mechanism of coding efficiency, offering deeper insights into neural coding strategies.

## 1 Introduction

One of the fundamental enigmas in the study of neural encoding revolves around the principles by which neurons encode sensory information, and the mechanisms they employ to effectively and dynamically represent ongoing information. Historically, the efficiency of neuronal coding systems has been tied to the number of neuronal response spikes, an aspect encapsulated by the concept of ‘response sparseness’ [1–3]. It suggests an efficient neural coding principle employing a few spikes from a few neurons in the network, organized sequentially and spatially, to represent various attributes of sensory inputs distinctively. This strategy highlights the brain’s remarkable efficiency [4]. Both experimental and theoretical studies have illuminated that in natural signal processing, salient features are often temporally represented through sequences of spikes that are sparsely distributed over time (lifetime sparseness) or spatially represented through the activation of a limited subset of neurons within a larger network (population sparseness) [1, 3].

Simultaneously, investigations of neural activities, both at the level of individual neurons and within neural populations recorded in vivo, have consistently exhibited high variability or irregularity [5, 6]. This variability was once considered a sign of intrinsic stochasticity [7], or a mark of the brain’s flexibility [8], adaptability [9], and coding capacity [10], and is quantitatively expressed through the ‘coefficient of variation’ (CV). CV, as a statistical metric, is computed as the ratio of the standard deviation to the mean of a specific distribution [11]. Depending on the particular application, it can be employed to gauge either the variability in the response of a neuronal population [12] or the variability over the lifetime of individual neurons [13].

The significant contrast between the high CV, indicative of inherent randomness in neural activity, and response sparseness, recognized as efficient neural information processing’s benchmark, necessitates a reevaluation of how neural systems reconcile the tension between randomness and operational efficiency. This dichotomy poses fundamental questions about the nature of neural processing: whether CV and sparseness operate as independent factors or converge on a yet-undiscovered unified mechanism, and how the metrics contribute to the nuances of neural coding and computation. Addressing these questions is crucial for unraveling the complexity of brain function and illuminating the underlying mechanisms of the notable efficiency and adaptability of the brain in processing information [14, 15].

Building on this framework, our research examines the fundamental link between CV-sparseness usages and their biological nature that advances neural computation. We begin by formulating an analytical equation that bridges these two metrics. Our findings suggest that the perceived ‘randomness’ of CV extends beyond simple stochasticity to represent a crucial aspect of energy-efficient coding, highlighting the deliberate utilization of irregular spike timings for efficient encoding information of input signals with minimal spike usage, thereby optimizing the energy efficiency of neural coding.

We further incorporate the two measures with a new metric, Information-Cost Efficiency (ICE), which encapsulates the coding capacity and the energy cost of neural responses. By proof of a positive correlation between CV and ICE, we show spike irregularity and response sparseness as a unified manifestation of an underlying strategy to maximize neural coding efficiency at a minimal cost.

To validate the above theoretical framework, we conduct simulations encompassing spike sequences from diverse inter-spike interval (ISI) distributions, a biophysical neuron model, and a spiking neural network exhibiting plasticity. These simulations reveal the neural mechanisms leading to increased sparsity or variability in neural responses invariably result in a corresponding increase in the other, highlighting the critical role of the excitation-inhibition balance in this reciprocity. This interplay has profound implications for understanding the dynamics of neural coding.

Furthermore, we develop a CV-maximization algorithm to learn natural images efficiently using the theoretical framework and successfully derive a comprehensive set of functional filters that closely mimic the receptive fields of the primary visual cortex. These results not only corroborate our theoretical findings but also provide a biological explanation for the creation of advanced, brain-inspired artificial neural network models and algorithms.

## 2 Theory

### 2.1 Theoretical Equivalence of CV and sparseness

Response sparseness and coefficient of variation converge to quantify neural coding efficiency. The sparse distribution and the high irregularity in the spike sequences are two properties that are widely and independently observed in the responses of individual neurons [16] and neuronal populations [17] in various regions of the brain. Historically, these properties have been quantified separately by response sparseness [2] and the CV [5, 6]. This study begins with exploring the equivalence of the measures to examine the potential for integrating them into a unified theoretical framework.

Here, based on experimental data[18] which includes simultaneous extracellular recordings from neuronal populations in the V1 visual areas of macaques, we observe the neuronal firing patterns in which the resting state shows a sparse firing in neurons, while the task state displays a more frequent firing. This shift in neuronal activity is further quantified through various spike train metrics presented in Figure 1A, B. In both patterns, the sparseness and CV are equivalent both in temporal and spatial space, see Figure 1C, E.

**Figure 1.**
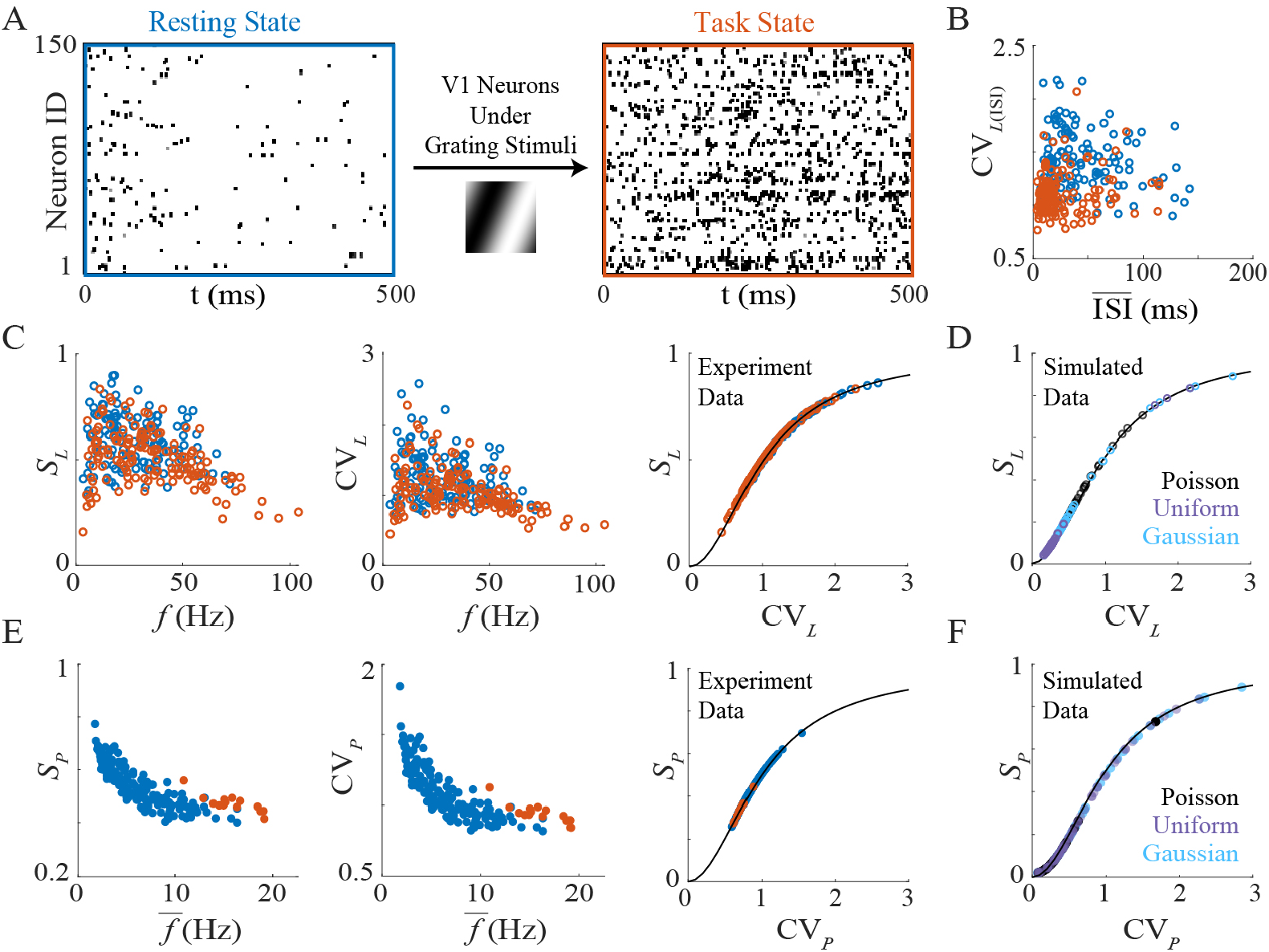
Sparse and irregular firing behaviors observed in single neurons and neuronal populations from experimental recordings can be quantified by both sparseness and the CV. (A) Rasterplot of neural spiking behaviors under resting and task state. (B) The high CV of interspike interals as suggested by [5]. (C) Lifetime sparseness (*S*_*L*_) and lifetime CV (CV_*L*_) of resting (blue circle) and task state recording (orange circle), together with (D) processes with the Gaussian (light blue circle), the Uniform (Violet circle) ISI distribution and Poisson process (black circle). (E) Population sparseness (*S*_*P*_) and CV_*P*_ of simulated processes and neuronal recordings (dots with corresponding colors as in C), together with (D) processes with the Gaussian (light blue dot), the Uniform (Violet dot) ISI distribution and Poisson process (black dot).

The concept of sparseness was initially introduced by Treves and Rolls [2], defined as a ratio:

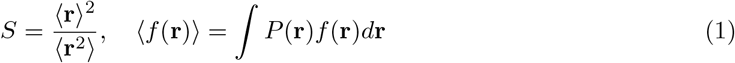

with *P* (**r**) indicating the probability density function of variable **r**. Vinje and Gallant later modified and extended this concept to capture the sparse firing patterns observed within neuronal networks in response to natural scenes [19]. For a set of neuronal firing rates **r** = *r*_1_, …, *r*_*N*_, for the temporal processes of individual neurons or population of neurons, the sparseness is revised and defined as [19]:

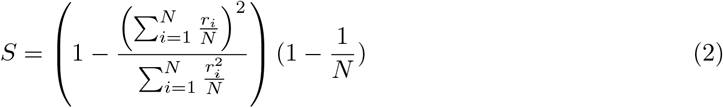

where *r*_*i*_ is the firing rates over time of an individual neuron or neurons in the network. Both the population [17] and lifetime [20, 21]sparseness reflect the energy efficiency of the input representa-tions. The terms 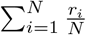 and 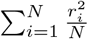 are the expectations of **r** and **r**^2^, denoted, respectively, by ⟨**r**⟩ and ⟨**r**^2^⟩. Thus, we rewrite the sparseness as follows:

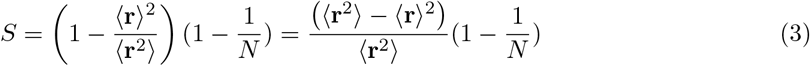

Then we prove that sparseness *S* is a monotonic increasing function of CV.

*Proof*. Notice that the variance *σ*^2^ can be written as:

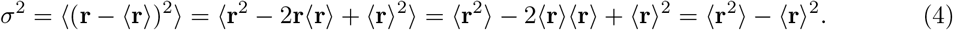

Then, we have:

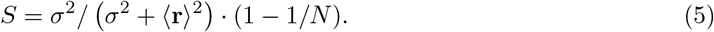

Recall that the neural response irregularity is quantified by *CV* = *σ/*⟨**r**⟩, then, we obtain

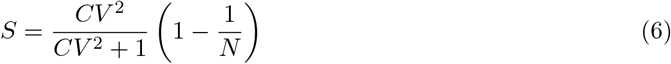

Given that the spike count N is generally large enough, i.e., (1 − 1*/N*) → 1, the expression can be simplified as:

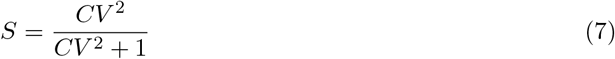

The derived equivalence equation proves that response sparseness *S* ∈ (0, 1) is a monotonic increasing function of *CV* ∈ (0, ∞) as long as the sparseness and the CV values are calculated to quantify the same response sequences of a given system (Fig. 1C-F, black lines), either along time or over network population. Both experimental data (Fig. 1C, E) and the simulation outcomes for spike sequences with various statistical distributions of ISIs (Fig. 1D,F) align perfectly with the predictions of the analytical curves (black line) calculated from Eq.7. The two measures of response sparseness and CV therefore have strong parallels. Note that the relationship between the two concepts was acknowledged in the field of statistics.

We further demonstrate that the equivalence between sparseness and CV holds for ISI distributions in Supplemental Materials(Figure S1).

Considering that the CV measure used in experimental studies is usually calculated over ISI [6, 12], hence, we also verify the monotonic relationship existing for lifetime *S*_*L*_ calculated by firing rate and CV_*L*(*ISI*)_ calculated over ISIs for Poisson sequences (Fig. S2, with a analytical proof in Supplemtal Proof 1.). Processes with Gaussian and Uniform ISI distributions are also validated to have a monotonic relationship between CV_*L*(*ISI*)_ and *S* (Fig. S3). This suggests a more direct relationship between sparseness and firing irregularity usually depicted by ISI for neuroscientists.

### 2.2 Integrate CV and sparseness into Information-Cost Efficiency

Efficiency quantification across various fields fundamentally involves optimizing performance while minimizing resource use. In energy efficiency, this optimization is measured by power consumption: Assessing computational output relative to energy costs [22]. For information coding efficiency, the principles of information theory guide maximization of information transfer [23]. When it comes to neural information processing systems, given the finite nature of energy resources, it is assumed that biological sparse encoding strategies represent an evolutionary adaptation within cortical circuits of animals [24, 25], necessitating a trade-off between maximizing functional output (high information capacity) and minimizing energy costs, as illustrated in linear models of predictive lateral coding [26].

To quantify such a trade-off, we consider entropy as a critical measure for quantifying the information capacity of neural systems, with a higher entropy value signifying an enhanced coding ability, suggesting more efficient utilization of resources in processing information [27, 28]. Conversely, energy expenditure is tied to the metabolic costs inherent to neural activities, encompassing the energy demands of synaptic transmission, neuronal firing, and maintenance of neural infrastructure.

We introduce a mathematical metric, **information-cost efficiency (ICE)** that synthesizes the two ends, aimed at measuring efficiency comprehensively. ICE is defined as the ratio of coding capacity, quantified by entropy in the context of information theory, over energy expenditure:

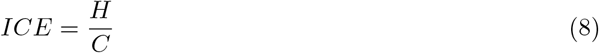

where *H* represents the information encoding capacity (measured by entropy), and *C* represents the energy of the temporal sequence itself. For simplicity, this analysis omits the additional energy costs associated with the coding hardware system [29, 30].

Taking into account a general signal {*x*_*i*_} that follows a distribution with mean value *µ* and standard deviation *σ*, CV is defined as *CV* = *σ/µ*. The power cost *C* of the given signal follows the definition of power [31] and the information capacity *H* [28] quantified by the entropy of a discrete statistical distribution are defined as:

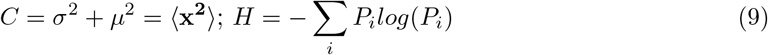

where *P*_*i*_ is the discrete probability of the distribution. The definition of *H* and *C* can be slightly modified considering the specific context.

Considering a set of variables that conforms to Poisson distribution:

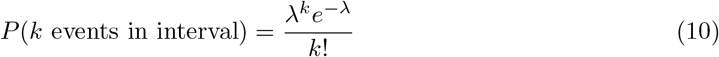

where *λ* is the average number of events per interval, *e* is Euler’s number (the base of natural logarithms), and *k*! is the factorial of *k*. Then, naturally, we have *µ* = *λ* and 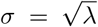, thus 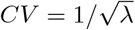.

Then we will have CV and ICE positively correlated. Here, we offer a proof for this positive relation:

*Proof*. The entropy and cost can be calculated respectively as:

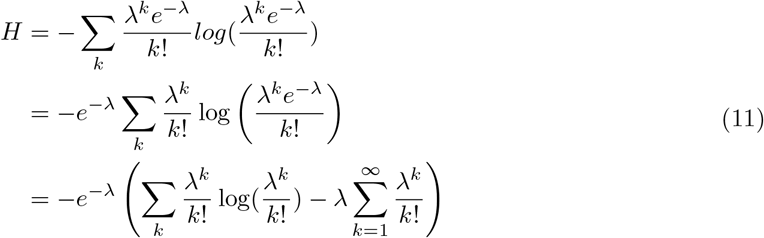

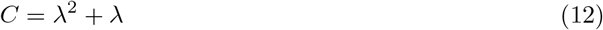

Then ICE can be written as a function *f* (*λ*):

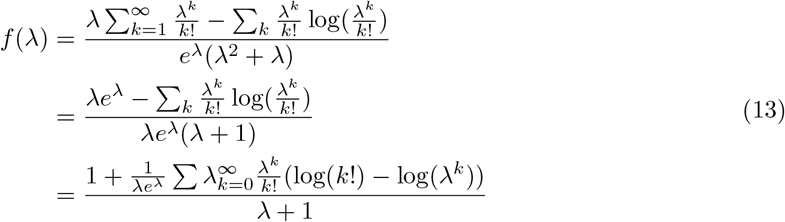

We have the derivative of *f* (*λ*) as:

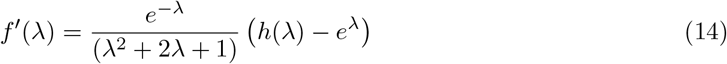

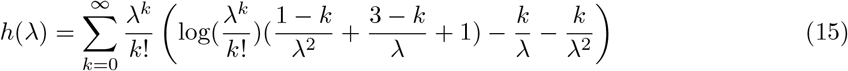

Since *h*(*λ*) is numerically validated smaller than *e*^*λ*^ on *λ* ∈ (0, 100) as suggested in Fig. S3, we have *f* ^*′*^(*λ*) *<* 0. If we denote *CV* = *x*, then *λ* = 1*/x*^2^, then:

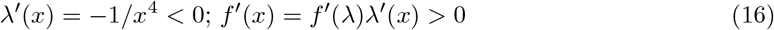

Therefore, we have *f* (*x*) as a monotonically increasing function of *x* ∈ (0.1, ∞). ICE is tightly correlated with CV for variables conforming to a Poisson process, suggesting a high CV indicating high Information-Cost Efficiency.

For entropy (*H*) and energy cost(*C*), we have:

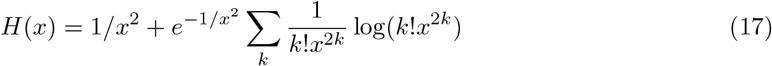

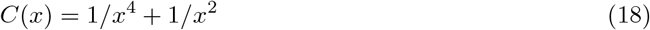

*ICE* = *H/C* is a function *f* (*x*) that can be written as:

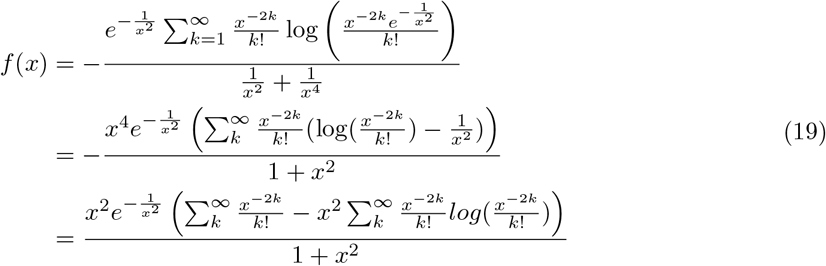

Thus, analytical relationships are derived between CV and key metrics such as entropy (Eq.17), energy cost (Eq.18), and information-cost efficiency (Eq.19), illustrated with black lines in Fig. 3. Additionally, ICE is analytically calculated for other statistical distributions, including Gaussian and Uniform distributions (see Supplemental Proof 3), where the mean values across distributions are adjusted to alter CV and, correspondingly, ICE (see Fig. S4). The correlation is also validated with the E-I network simulation (see Fig. S5, details in Materials and Methods 5.4).

**Figure 2.**
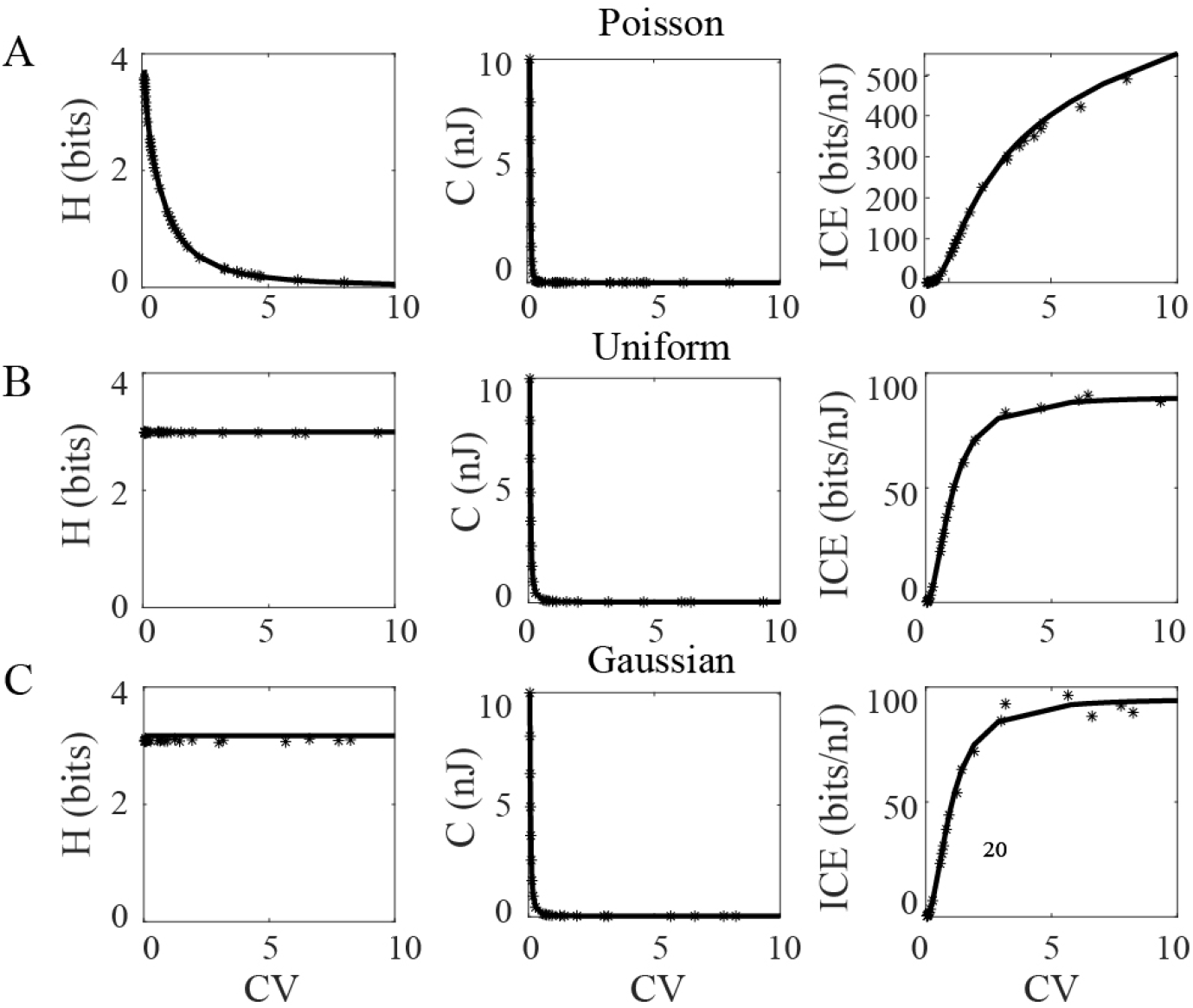
ICE is a monotonically increasing function of CV for variables that conform to Poisson distribution given the analytical and numerical evidence. Entropy (*H*) and Cost (*C*) plot with CV of (A) Poisson distribution, (B) Uniform distribution and (C) Gaussian distribution.

**Figure 3.**
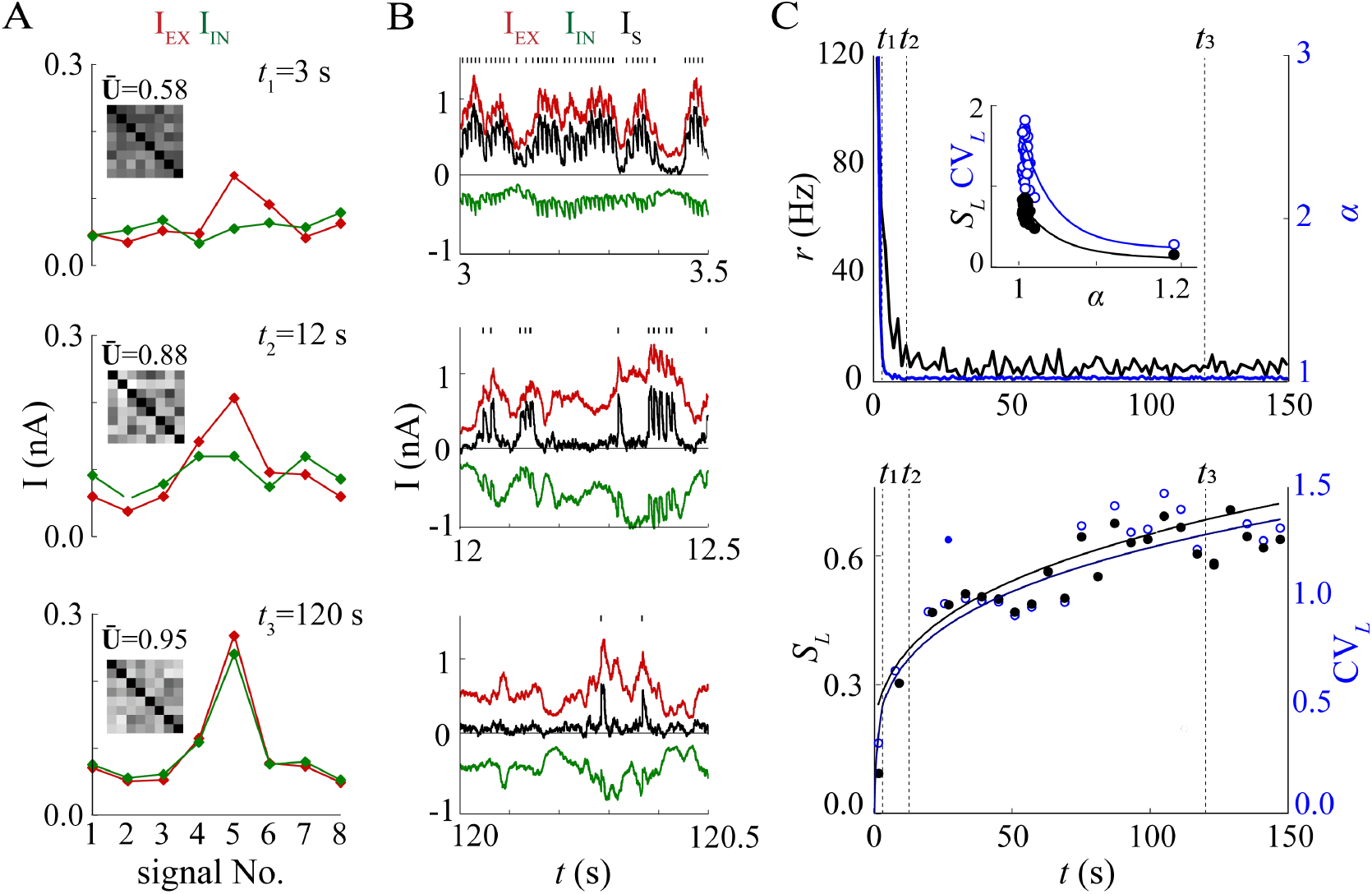
Synaptic plasticity enhances neuronal spiking variability and lifetime sparseness. (A) The mean amplitude of the excitatory (*I*_*EX*_, red) and inhibitory (*I*_*IN*_, green) currents for eight input signals in a 1.5-second time window starts at *t*_1_, *t*_2_, and *t*_3_ (*t*_1_ = 3*s, t*_2_ = 12*s*, and *t*_3_ = 120*s*, marked in panel C by three dashed lines). The dissimilarity of each pair of signals is suggested by the matrix with mean values *Ū* illustrated below. (B) Total excitatory and inhibitory membrane currents (*I*_*EX*_, red / *I*_*IN*_, green) and sum (*I*_*S*_, black) over time in a time window of 500 ms before (*t*_1_), during (*t*_2_), and after (*t*_3_) implementation of synaptic plasticity. Corresponding spikes were recorded above each plot. (C) The firing rate decreases over time and high values of CV_*L*_ and *S*_*L*_ are obtained when *alpha* is close to 1 (top), CV_*L*_ and *S*_*L*_ (bottom) as functions of time.

## 3 Results

### 3.1 E-I balance underlying high CV-sparseness and ICE of neural activities

Our previous studies [21, 32, 33] as well as other experimental [13, 34] and computational studies [35] over the past decades have established several key neural mechanisms to achieve large response sparseness or CV in neuronal networks during processing sensory inputs. The high CV in neural activity is shown to not be due to the intrinsic Gaussian-type noise distribution but is most significantly determined by the concurrent balanced ratio developed between excitatory-inhibitory synaptic inputs [33, 36] during the sensory learning process, as well as the intrinsic excitability properties of the neuronal membrane [37, 38].

In this section, we conduct simulations of general models on biophysical neurons and spiking neural networks (models are adapted from previous studies [19]) to explore the common mechanisms behind the high CV-sparseness and the energy efficiency of neural activities. These models offer profound insights into synaptic plasticity’s role in achieving the balance between excitation and inhibition within neural circuits, leading to irregular asynchronous network states. These highly asynchronous activities are crucial for the effective functioning of sensory pathways and memory networks [19]. We will demonstrate that high CV-sparseness is a characteristic outcome of a stable, low-cost dynamic state, emerging from the optimal balance process between excitation and inhibition. This enables the target neuron to more effectively distinguish between sets of input signals through unique irregular and sparse temporal spike sequences, propelling the network towards a spatially asynchronous state that attains higher information-cost efficiency.

#### 3.1.1 Balanced E-I synaptic input leads to temporally sparse and irregular firing of individual neuron

The target neuron receiving concurrent excitatory and inhibitory synaptic input exhibits large CV_*L*_ and *S*_*L*_ when timing and strength of excitatory and inhibitory synaptic input are tightly balanced after the long-term learning processes of eight given signals (Fig. 3, see Materials and Methods 5.3 for details). As the input signal amplitudes and membrane currents of a target neuron become more balanced (E/I balance ratio *α* = *I*_*EX*_*/I*_*IN*_ approaches 1) before, during, and after the implementation of inhibitory synaptic plasticity (Fig. 3A, 3B), the target neuron gradually obtains maximal values of *S*_*L*_ and CV_*L*_(Fig. 3C). The pairwise dissimilarity matrix *U*_8*×*8_ of the input signals is calculated in the corresponding time windows, with its upper triangle representing the pair dissimilarity of the excitatory signals Nos. 1-8 and the lower triangle represents inhibitory signals. The increase in the mean value of the dissimilarity matrix *U* suggests an improved ability to distinguish the 8 groups of signals when the target neuron spikes more sparsely. The instantaneous firing rates are recorded to calculate the *S*_*L*_ and CV_*L*_ values for every 6-second time window. The growing values certify that balanced excitatory-inhibitory synaptic currents are indeed the underlying dynamical mechanism of high sparsity and large irregularity.

#### 3.1.2 Self-evolving E-I balance leads to spatially sparse, irregular, and asynchronous network activities

To further examine the role of E-I balance for efficient spatial patterns of a neuron population, we simulate a spiking neural network (see Materials and Methods 5.4 for details) that encompasses both excitatory and inhibitory neurons, interconnected through synaptic links governed by Hebbian plasticity. This network shows spatially sparse and irregular activation of a limited subset of neurons after implementing inhibitory plasticity at t = 4 s. The mean firing rates of both excitatory and inhibitory groups decrease rapidly during the adaptation process. The E/I ratio *α* (calculated by the mean firing rate of the excitatory neuron group over the inhibitory group) rapidly decreases to 1, leading to growing values CV_*P*_ and *S*_*P*_ that perfectly matched the CV-sparseness equation (Fig. 4E-F). This process is accompanied by a transition from a quasi-synchronous state to the emergence of a globally asynchronous network state when excitation and inhibition of a network reach a tightly balanced level.

**Figure 4.**
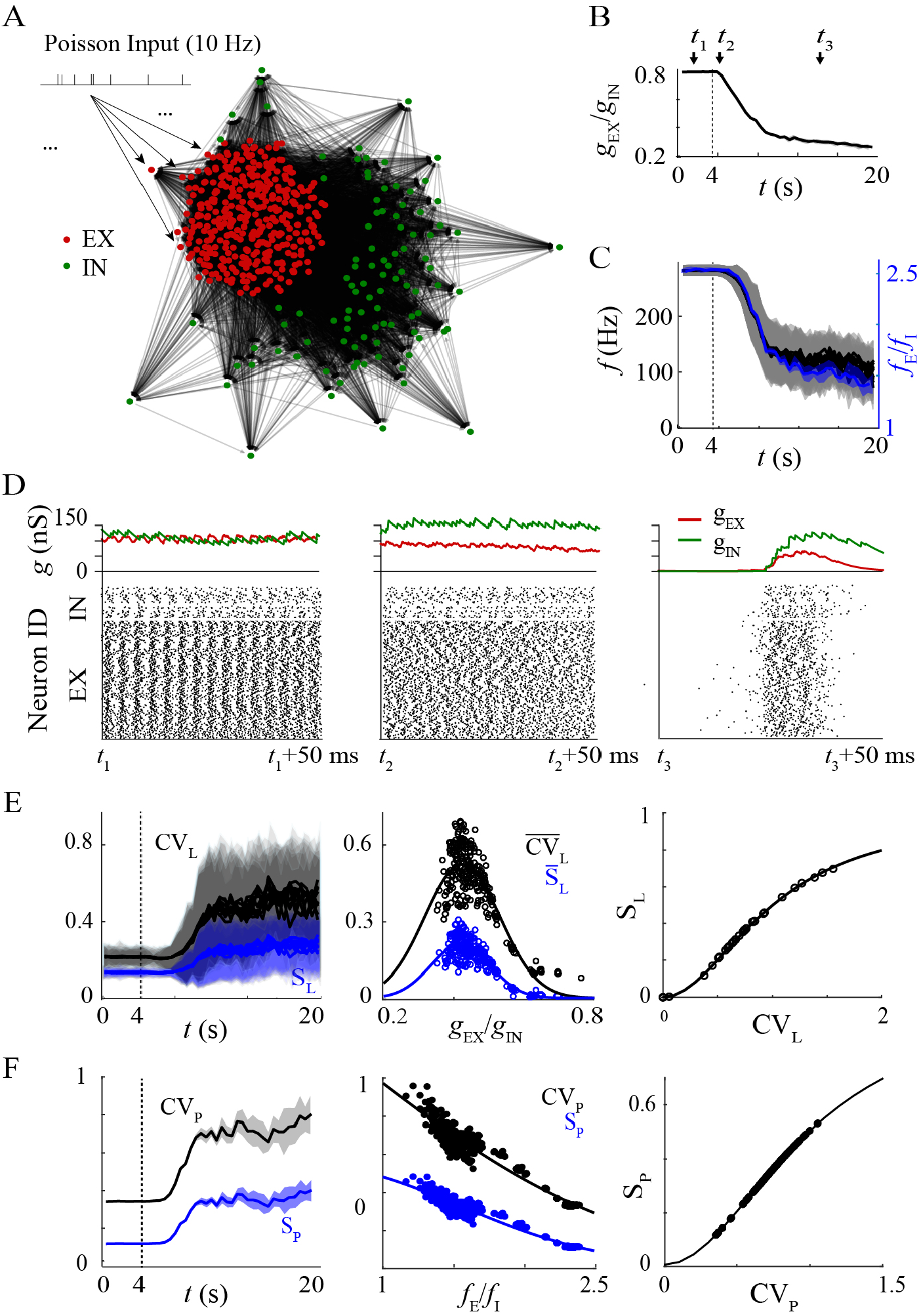
Population sparseness and firing variability of a network over time. (A) Illustration of the E-I network. After implementing inhibitory plasticity at *t* = 4 s: (B) the ratio of excitatory and inhibitory synaptic strength decreases while (C) the mean firing rate of both the excitatory and inhibitory neurons and E/I ratio *f*_*E*_*/f*_*I*_ decreased. (D) 50 ms spike raster plots and the excitatory (red) and inhibitory (green) synaptic inputs to each neuron before (*t*_1_), during (*t*_2_), and after (*t*_3_) employing inhibitory plasticity (Note that *t*_1_, *t*_2_, and *t*_3_ are marked in Panel A and B by the 3 arrows). (E) The *S*_*L*_ and CV_*L*_ values of the excitatory and inhibitory neurons both increased to a stable level. The peak reached with a specific ratio of synaptic E-I balance. The data fit well with the CV-sparseness relationship. (F) The *S*_*P*_ and CV_*P*_ values of the excitatory and inhibitory neurons both increased to a stable level. Larger CV and sparseness reached with less excitatory activities E-I balance. The data also fit well with the CV-sparseness relationship.

#### 3.1.3 Achieving augmented ICE along with enhanced CV-sparseness through E-I balance

Now We are investigating the impact of growing CV-sparseness on neural coding efficiency by utilizing the Information-Cost Efficiency (ICE) metric. This metric allows us to assess the balance between information encoding and energy consumption within the network. Our analysis highlights the significance of maintaining balanced excitatory and inhibitory interactions for efficient coding.

As depicted in Fig.4D, the network transitions from a synchronous to an asynchronous state during the plasticity process. This ultimately leads to a balanced excitation-inhibition state, as shown in Fig. 4. Interestingly, during this transition, we observe a concurrent increase in the average information *H* per neuron (Fig. 5A) and a decrease in the average firing rate and associated energy cost (Fig. 5D). This translates to a rise in the ICE metric (Fig. 5G), indicating enhanced energy-efficient coding within the network.

**Figure 5.**
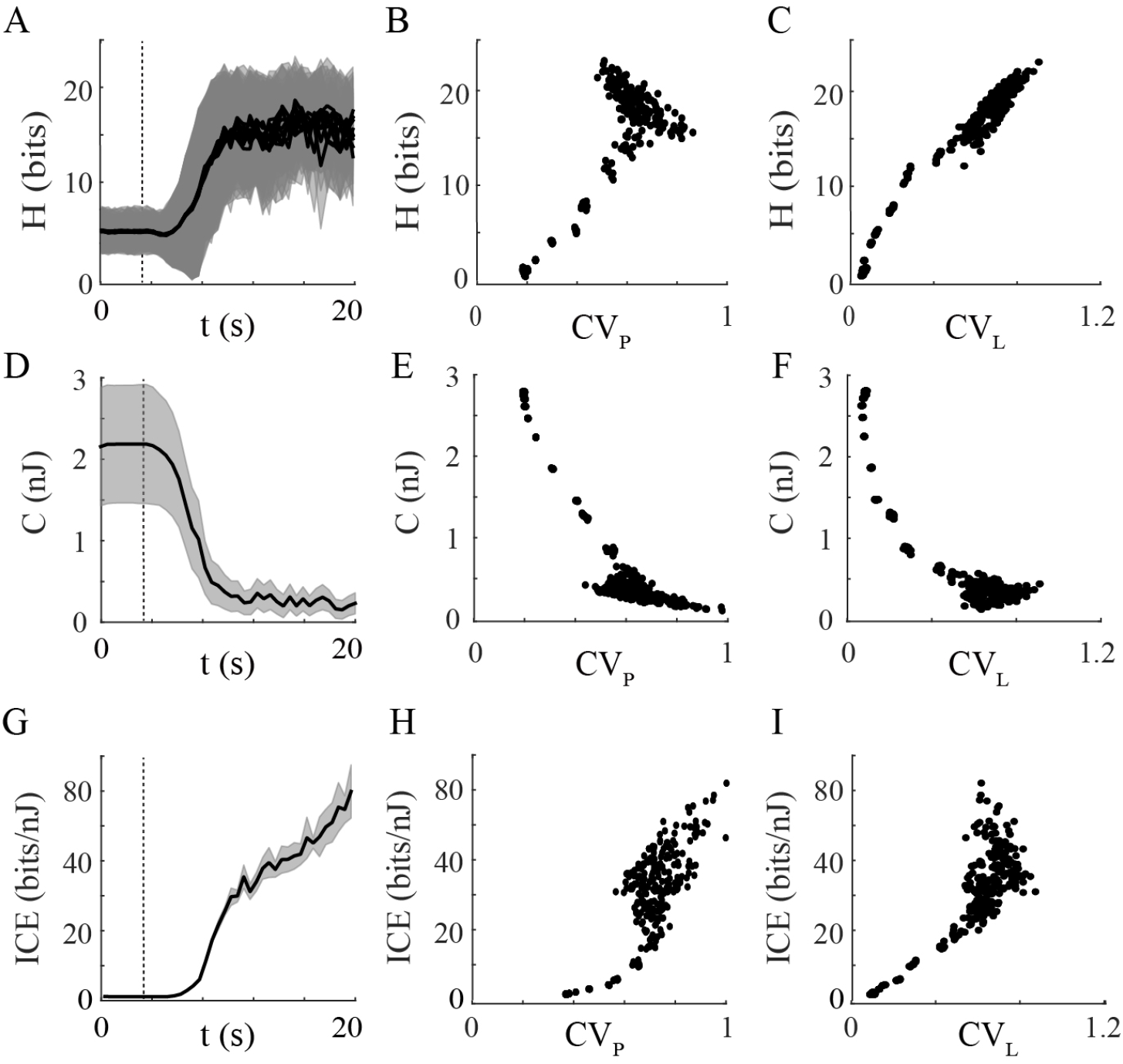
The E-I neural network demonstrate temporal increases in the Information-Cost Efficiency (ICE) metric. In the E-I network (same as shown in Fig. 4), the average entropy *H* per neuron increases as the system transitions from a synchronous to an asynchronous state (A). Postive relation between *H* and (B)*CV*_*P*_, (C)*CV*_*L*_ both observed during adaptation. Meanwhile, the introduction of plasticity leads to a decrease in energy cost *C* (D-F). This combined effect results in an improved ICE value (G-I), indicating improved coding efficiency. Mean values across trials or neurons are drawn by black lines with standard deviation in shades.

In sum, with the proven correlation between ICE and CV-sparseness, we conclude that enhanced response sparseness and CV are linked to lower energy consumption due to decreased firing rates, thus leading to higher ICE values signifying substantial information capacity within constrained energy budgets for a system [39].

### 3.2 CV maximization algorithm for deriving complete basis functions of visual receptive fields

Biological visual receptive fields are crucial for salient feature extraction from visual scenes, with each neuron selectively responding to specific principal components within images, ensuring efficient image encoding [40, 41] while different neurons have distinct receptive fields that are specifically designed to detect different characteristics of the visual environment [42], reducing redundancy and allowing the brain to represent visual signal more efficiently. Furthermore, the bio-receptive fields are observed to adapt to the statistics of visual environment [43] and experience [44], and this adaptability is essential to optimize visual processing in a changing environment.

Biological vision systems, despite their complex processing, achieve remarkable energy efficiency. Neurons fire only as needed, with receptive fields selectively responding to relevant stimuli, minimizing energy use on non-essential information [45]. This energy-conserving mechanism, along with their ability to extract salient features efficiently, positions biological visual systems as models of efficiency and specialization, offering valuable insights for designing energy-efficient artificial systems.

Having confirmed with the above results the potential of nonlinear coding systems to efficiently encode signals, suggesting that further exploration of high CV-sparseness could bridge the efficiency gap between biological neural systems and contemporary artificial neural networks, we now consider the functional significance of the sparse and irregular nature of neural systems and how it can help to develop advanced image processing algorithms mimicking the biological visual system along with similar receptive fields.

Inspired by the prominent advantage of response sparsity and efficient coding of neuronal systems, the sparse coding algorithm addressed has been proven to be powerful in machine learning [46– 48]. By using a sparse coding strategy that maximizes the sparseness of weight for the set of basis functions, Olshausen and Field successfully developed a complete family of localized, oriented, bandpass receptive fields, similar to known neuronal functions in the primary visual cortex [49]. As the complexness of the mathematical definition of sparseness has limited its application, the established equivalence between CV and sparseness here allows us to introduce a new training algorithm that maximizes CV of basis weights, replacing the original sparse-coding framework.

The concept of sparse coding brought up by Olshausen and Field [49] is an unsupervised learning method used to find a set of “overcomplete” basis vectors Φ(*x, y*), *i* = 1, …, *N* to efficiently represent the input image *I*(*x, y*) as a linear combination (weighted sum by coefficient *a*_*i*_, *i* = 1, …, *N*) of these basis vectors. Trained with small patches randomly selected from natural images, the network generated 144 bases (N=144), each consisting of 8×8 voxels. Then given a test patch, we can reconstruct the patch with a linear combination of basis as in Fig. 6A, and the coefficient, i.e. the weight vector, is sparse (there exist only few non-zero terms in a vector of 64 dimensions). For the null model, we directly train the network minimizing the loss function *L*:

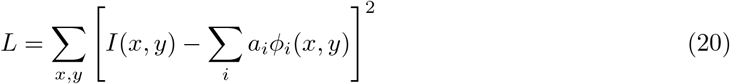

**Figure 6.**
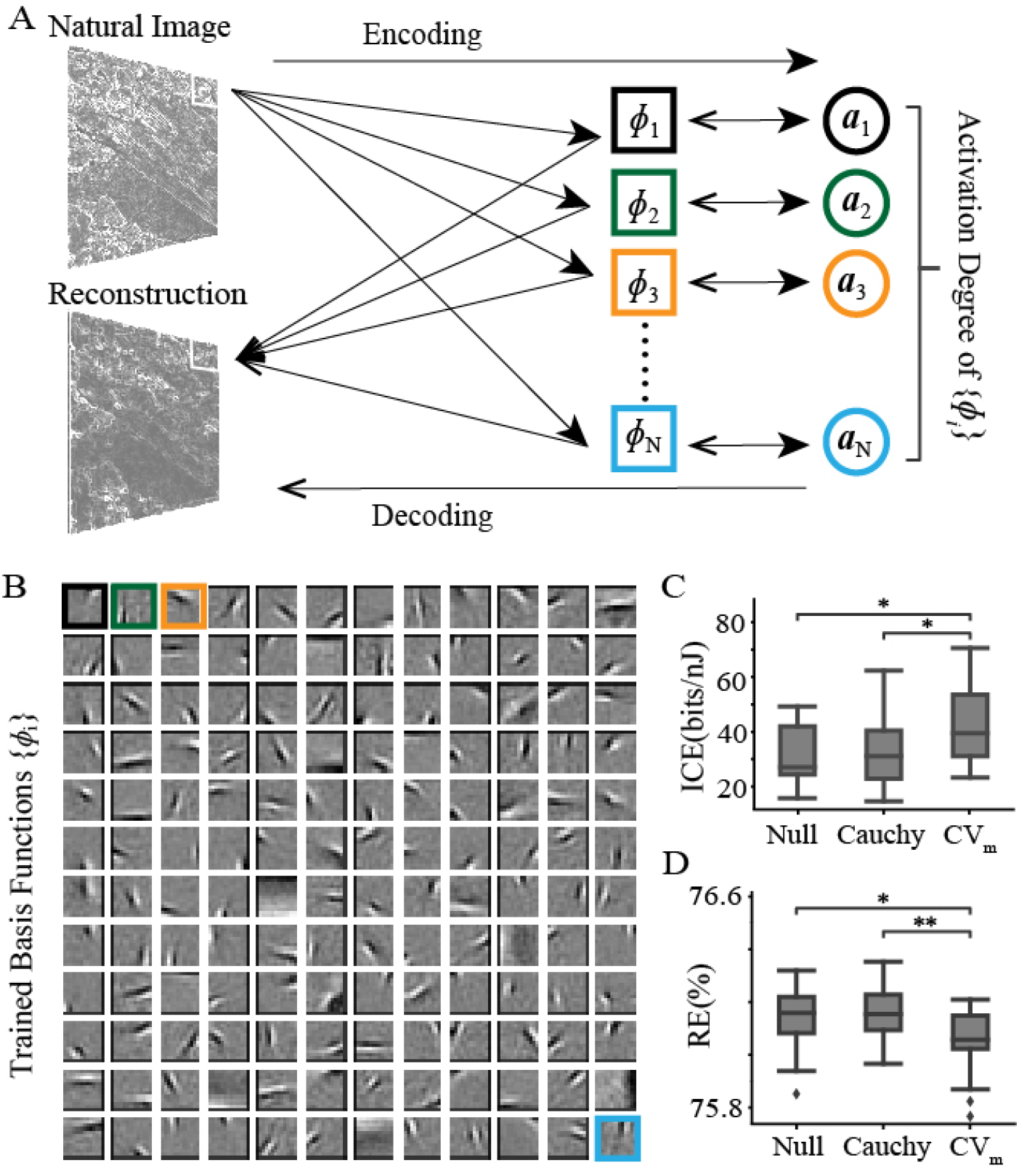
(A) Maximizing the coefficient of variation during network training on 16×16-pixel image patches from natural scenes results in a set of learned basis functions. These functions closely mimic the characteristics of the receptive fields found in the primary visual cortex of mammals. *a*_*j*_, *j* = 1, …, *M* are a set of N-dimensional vectors that suggest weights of N basis functions while reconstructing an image of M patches. Basis functions are illustrated in (B). The boxplots compare the performance of three distinct approaches: a null model focused solely on minimizing reconstruction error, an approach that applies the Cauchy prior constraint as outlined in the sparse coding algorithm [38], and a method that aims to maximize CV. Performance is illustrated by (C) ICE and (D) reconstruction error (RE). Of these metrics, the CV maximization method demonstrates superior performance in terms of accuracy compared to both the null model and the original sparsecoding algorithm.

In previous study[49] constrained by sparse coding principle, the weight vector is assumed to conform to a parametric prior distribution:

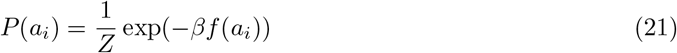

where *f* (*a*_*i*_) is a nonlinear function that determines the shape of the prior distribution, and a Cauchy distribution *f* (*a*) = *log*(1 + *a*^2^) was used by Olshausen and Field [50], *Z* and *β* are scaling constants. Since each weight vector a is to be regularized to obtain a sparse distribution, the loss function is to minimize the reconstruction error and nonlinearity of weights, defined as:

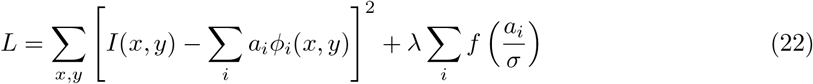

Here, *σ* and *λ* are also scaling constants. The absolute value of weight can be regarded as a coupling of connection strength and firing of a neuron, then sparseness of weight will naturally correlate to sparseness of neural activities biologically.

However, defining biological sparseness as outlined in Eq.2 involves complex calculations. Given the derived “CV-sparseness” equivalence, we propose a new algorithm that focuses on maximizing the CV of the weight distribution during training, rather than directly targeting sparseness. This approach eliminates the need for assuming a prior distribution:

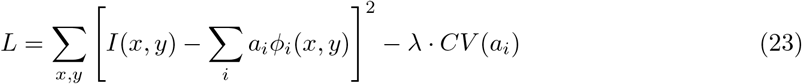

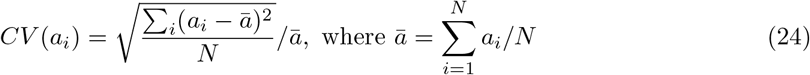

As shown in Fig. 6, maximizing CV leads to a full set of filters with localized, oriented, bandpass features, akin to the primary visual cortex’s receptive fields. Using these filters for input image reconstruction results in higher performance, with lower reconstruction error (RE) (calculated as Eq.20) and higher ICE than both the null model and the sparse-coding model with Cauchy prior. Our model optimizes weight distribution through loss minimization with CV-maximization constraint, while the original sparse-coding model chooses what seems to be the best probability distribution based on experience, which does not achieve the best theoretical results. Thus the filters derived by Cauchy prior still produce noticeable errors in image reconstruction. Structural similarity index (SSIM) and dissimilarity among the derived basis (measured by the mean value of *U*_*ij*_ = 1 − *corrcoef* (Φ_*i*_, Φ_*j*_)) are also compared as in Fig. S6. The learning processes were repeated in 20 trials.

Maximizing CV of weight distribution results in filters with statistically independent basis that resemble visual receptive fields, leading to superior image reconstruction and higher coding efficiency. Thus, the CV-maximizing algorithm offers a robust framework for machine learning which may deepen our understanding of neural coding principles.

## 4 Discussion

### 4.1 Unified Framework Reconciling Sparseness and CV with ICE

In this study, we present a unified theoretical framework that deepens our understanding of neural encoding by elucidating the profound interconnections between coefficient variation (CV) and sparsity in neuronal responses. This framework posits that the mechanisms driving sparsity [20, 21, 33] or variability [32, 51–53], particularly the crucial balance between excitation and inhibition (E-I balance) [3, 27, 54, 55]—an essential factor in neural function—are fundamentally intertwined, profoundly influencing both attributes. It not only aligns these properties with the principles of information-cost efficiency (ICE) but also resolves enduring debates over the functional implications of neuronal variability and sparsity. By systematically examining the interplay between these metrics, our analysis underscores their essential roles in enhancing neural efficiency through a balanced economy of information processing and energy conservation.

Historically, the variability observed in neuronal spiking has been a contentious topic, with debates centered around whether it represents inherent stochasticity, potentially detrimental to neural information processing [7], or serves as a substrate for enhanced coding capacity [28, 56, 57]. Our findings bridge this divide by demonstrating that such variability is far from merely stochastic; it is indicative of a highly efficient coding strategy utilized by biological brain circuits, especially when analyzed through the ICE framework. This insight provides a crucial piece of the puzzle in understanding the adaptability and efficiency of biological neural networks, suggesting that neuronal irregularity is not meaningless randomness but a fundamental aspect of energy-efficient neural computation.

Furthermore, the integration of sparsity and CV within our theoretical framework offers a transformative perspective on the dynamics governing neural activity. By demonstrating how these metrics, traditionally viewed as distinct or competing, actually function as complementary forces, our approach challenges conventional beliefs and illustrates their synergistic evolution to enhance the brain’s information-processing capabilities. This paradigm shift, underpinned by the principles of entropy and cost from the field of information processing [58, 59], not only enriches our understanding of neural coding dynamics but also pioneers the development of more sophisticated computational models. These models emulate the brain’s energy-efficient strategies, optimizing both the transmission and processing of information by fine-tuning neuronal activation patterns. This results not merely in enhanced information efficiency but also in significant reductions in energy consumption, laying the groundwork for the creation of bio-inspired algorithms that effectively balance system disruption with the benefits of cost efficiency.

### 4.2 Correspondence with Free Energy Principle

The mathematical nature of CV, as we explained with the ICE framework, has suggested an intrinsic trade-off between information capacity and energy cost that can reveal the cost efficiency of the neural system. This can also be understood from the perspective of the free energy principle. In physics and physical chemistry, free energy is a measure of the reversible work that a system can perform under a given temperature and pressure, and it is a key bridge between the energy and entropy of the system. The concept of free energy occupies a central place in thermodynamics and statistical physics and is used to describe the change in energy of a system as it reaches thermodynamic equilibrium [60].

The ICE index quantifies the efficiency of information encoding and transmission of the nervous system per unit of energy expenditure, which is consistent with minimizing the free energy of the system to achieve optimal information processing in the free energy principle [61]. Considering the format of the 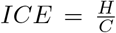 and expression of free energy *F* = *U* − *TS*, we can observe consistent ideas of minimizing free energy *F* and energy cost *C* while maximizing entropy *H* or *S* of a system given certain internal energy of the system *U*. Thus, the self-evolving process of the neural network during neural coding, as it moves toward a state of maximal energy efficiency, may indicate the achievement of minimal free energy and maximal entropy. This aligns with the free energy principle as the system nears thermodynamic equilibrium. Associating the Information-Cost Efficiency (ICE) metric with the free energy principle offers a novel perspective on the economics of energy-efficient neural coding, interpreted through the principles of physical science.

In the context of biology, the free energy principle has been introduced to neuroscience by Karl Friston [62] to emphasize the optimal behavior of biological systems in perception and action, enabling adaptation to the environment by reducing prediction errors with defined variational free energy. However, this application neglected the energy consumption of biological processes to some extent. Comprehending the ICE indicator with the original definition of free energy evaluates the energy utilization considering not only the accurate transmission of information but also the realistic energy conservation of the system, which is important for understanding how the brain achieves complex cognitive functions with a limited energy budget.

### 4.3 Implications for novel AI algorithms

Leveraging the ICE metric from the CV-sparseness relationship, we offer an analytical lens to explore brain optimization strategies, further clarify the role of CV-sparseness in neural coding in existing knowledge [63], and emphasize the efficiency of sparse spiking in neural coding, which is now widely employed in ANN designs [64]. This framework for analyzing efficiency in both natural and artificial neural networks is vital for creating artificial neural networks that emulate biological neural efficiency [65].

Incorporating strategies to improve sparseness and CV, mirroring natural optimization, significantly enhances ANNs in computational efficiency and performance. This approach is particularly crucial in resource-constrained scenarios [66, 67] for boosting ANNs’ efficiency while supporting sustainable computing. However, it is essential to heed previous warnings that solely maximizing spiking variability or sparsity might compromise the fidelity of neural coding [68]. This suggests a sophisticated optimization within cortical circuits, striking an ideal balance between encoding capacity and fidelity, all while adhering to energy constraints. In the future, we can include this balance, which is crucial for the dependability of neural processing systems, into the ICE framework and sets the stage for future research aimed at underscoring the feasibility of energy-efficient but also highly reliable computational models.

### 4.4 Towards biologically interpretable efficient information processing systems

Generalizing the CV-sparseness framework to various information processing systems is promising, given the ubiquity of large CV across disciplines, especially in biostatistics [69, 70], suggesting its foundational role in efficiency and reliability in broader information processing contexts [71]. Here, our newly developed CV-based feature-identification algorithm maximizes weight distribution’s CV, creates filters akin to the visual cortex’s receptive fields, and surpasses traditional sparse coding in image reconstruction, base dissimilarity, and ICE. Extending the CV-sparseness framework to a broad spectrum of information processing systems could foster an integrated approach to studying efficiency and adaptability, leading to computational models that mimic the natural balance between sparsity, variability, and performance [72]. This interdisciplinary effort could bridge fields from artificial intelligence to biotechnology, clarifying fundamental principles and catalyzing innovations that leverage efficiency and adaptability in complex systems [73].

By establishing the equivalence between sparseness and CV, and further integrating this understanding with the ICE metric, we provide a solid biological foundation for obtaining highly efficient information processing systems by employing CV-related metrics. With interpretability becoming an arousing topic in the field of ANN [74, 75], this research demonstrates the transformative impact of cross-disciplinary endeavors on deepening our understanding of neural coding’s efficiency and thus enhances the biological interpretability for current neural network models optimized with statistical metrics. Thus, the derived framework provides future directions that have the potential to greatly promote our grasp of brain functionality and propel the development of computational models that emulate the remarkable efficiency of biological systems.

### 4.5 Conclusion

In this investigation, we present a groundbreaking theoretical framework that resolves a foundational challenge in neuroscience—enhancing our understanding of how neural systems achieve highly efficient information processing with constrained energy resources. By establishing a mathematical link between coefficient variation (CV) and response sparsity, and integrating these with the novel metric of information-cost efficiency (ICE), our study provides a transformative perspective on neural coding. This approach not only offers a rigorous theoretical underpinning for optimizing neural activity but also elucidates the sophisticated computational strategies inherent in biological neural networks. These results underscore the remarkable adaptability and efficiency of biological systems, particularly the human brain, in managing vast quantities of sensory information while minimizing energy consumption. This efficiency is not merely a characteristic of neural operation but a fundamental aspect of neural evolution and function. The implications of our findings extend beyond neuroscience, offering valuable insights for the design of novel, biologically inspired computational architectures. These models promise significant advancements in energy efficiency and performance, particularly in environments where resources are limited.

Moreover, this study has the potential to influence the fields of artificial intelligence and machine learning. By applying the principles uncovered through our research to artificial systems, we can advance the development of more intelligent, efficient, and sustainable computing technologies. Such technologies are crucial for the next generation of AI applications that require lower energy consumption while maintaining high performance.

In conclusion, by effectively mimicking the brain’s computational strategies for efficient coding, we open new avenues for enhancing computational systems. These advancements are poised to make substantial impacts across various scientific and engineering disciplines, reinforcing the role of neuro-inspired models in the ongoing evolution of technology.

## 5 Materials and Methods

### 5.1 Extracellular recordings of V1 neurons in macaque brain

The experimental dataset discussed [18] originates from a subset of data reported in [76], captured using a sophisticated 96-channel Utah array with specific dimensions and depth, aimed at recording V1 activity. Neurons sampled for this study were located close to the fovea in the lower visual field, with their receptive field sizes quantitatively defined by fitting a Gaussian function to the data. The study employed drifting sinusoidal gratings as visual stimuli, presented in eight distinct orientations to investigate the responsiveness of V1 neurons. Each stimulus lasts for 1.28 s, followed by a blank screen for 1.5 s. Spikes were counted in 100 ms bins starting 160 ms post-stimulus onset, with the analysis period extending over one second per trial.

### 5.2 Neuronal spike sequence simulation with different ISI distributions

To numerically investigate the relationship between CV and sparsity, we are conducting a series of simulations using generated neuronal spiking sequences based on Poisson processes (depicted as cyan circles in Fig. 1, black/dark/light gray dots in Fig. 2), Gaussian (represented by red circles in Fig. 1), and uniform (indicated by blue circles in Fig. 1) interspike interval (ISI) distributions. Each simulation is being assessed using both *S*_*L*_ and CV_*L*_, which are being calculated over the firing rate within each 500ms time window of the simulated sequences lasting 50s, as shown in Fig. 2. Furthermore, both *S*_*P*_ and CV_*P*_ are calculated over the mean firing rate of each Poisson sequence in Fig. 2. The mean firing rate ranges from 2 to 100 Hz, while an absolute refractory period of 1 ms is maintained for all the simulated processes in Fig. 1. (Refractory periods of 2 ms and 3 ms are being tested for Poisson processes, with the results illustrated in Fig. 2).

### 5.3 Single neuron model learns to discriminate eight distinct signals

We build a working memory model system that features a target LIF neuron that receives 8 independent traces of low-pass filtered, half-wave rectified white noise signals to mimic sensory input during learning, all represented in parallel through correlated excitatory and inhibitory currents from eight groups of neurons (100 excitatory neurons and 25 inhibitory neurons in each group).

The LIF model is used as follows:

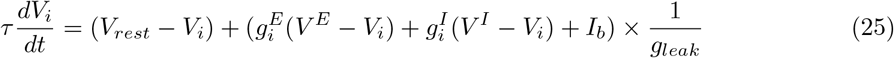

where reversal potentials are *V*_*rest*_ = −60 mV, *V*_*E*_ = 0 mV and *V*_*I*_ = −80 mV, *g*_*leak*_ = 10 nS, *τ* = 20 ms and *I*_*b*_ = 0 pA for single neuron model. The spiking threshold is -50 mV and an absolute refractory period of 5 ms exists after each spike.

Excitatory synapses were set at 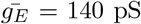, and the weight factor *W*_*ij*_ was adjusted to obey the (arbitrary) function *W*_*ij*_ = 0.3 + 1.1*/*(1 + (*K*(*j*) − *P*))^4^ + *ξ*, where *K* ∈ 1…8 is the group index of the presynaptic synapse *j, ξ* ∈ [0…0.1] is a noise term and *K* = 5 is the position of the peak of the tuning curve, that is, the signal with the strongest synapses. Inhibitory synaptic conductances were initially set to 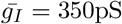.

This spike-timing dependent plasticity (STDP) rule was implemented for inhibitory synapses projecting onto excitatory cells [77, 78] with the synaptic trace:

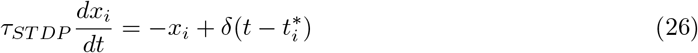

where 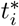 is when neuron i emitted a spike. Thus, the synaptic weight *W*_*ij*_ from neuron *j* to neuron *i* is updated for every pre-or postsynaptic event such that

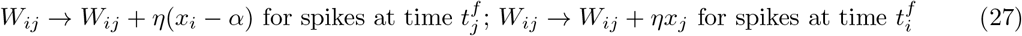

where *η* = 0.0012 is the learning rate and *α* = 2 × *ρ*_0_ × *τ*_*STDP*_ is the depression factor, *ρ*_0_ is a scaling constant parameter with units 1/time [19], here we set *ρ*_0_ = 5Hz, and *τ*_*ST DP*_ = 30 ms. 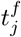 and 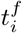 are time at which presynaptic spikes of the neuron *j* and postsynaptic spikes from neuron *i* occur accordingly.

### 5.4 The excitatory-inhibitory neural network employs inhibitory plasticity

To investigate the mechanism underlying the population sparsity and variability on neural networks, we built an excitation-inhibition LIF neural network with 100 inhibitory and 400 excitatory neurons that employ synaptic plasticity. Neuronal dynamics also follows Eq.25 with *I*_*b*_ = 100 pA and a refractory period of 3 ms. Within the neuron population, inhibitory plasticity is used to regulate the equilibrium between excitatory and inhibitory synaptic currents [19], the plasticity model is the same as in Eq.26 with *τ*_*ST DP*_ = 20 ms.

The synapse model follows:

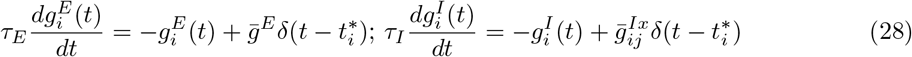

where *τ*_*E*_ = 5ms and *τ*_*I*_ = 10 ms, 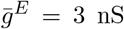. For 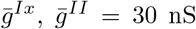, and 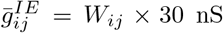, with *W*_*ij*_ ∈ (0, 30) follows the Eq.27. Here, for the network simulation, we set *η* = 0.001 and *α* = 2 × *ρ*_0_ × *τ*_*STDP*_ = 2 × 3 × 0.02 = 0.12.

The neurons are randomly connected with a probability of 0.12 from the excitatory group to both the excitatory and inhibitory groups and also a probability of 0.12 from the inhibitory to both groups. The values of *S*_*P*_ and CV_*P*_ are calculated based on all neuronal firing rates for each 1-s window.

### 5.5 Sparseness and CV calculation for neuron population and single neuron

The definition of the population/lifetime sparseness *S*_*P*_ /*S*_*L*_ and the CV_*P*_ /CV_*L*_ are correspondingly based on the mean firing rate of each neuron in the population and the mean firing rate in windows for a single neuron over time. We can calculate with:

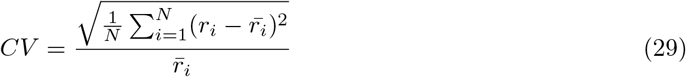

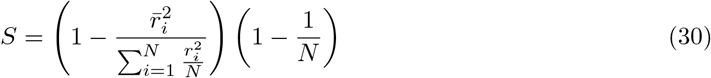

where 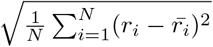 is the standard deviation of the firing rate, while 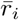 is the averaged firing rate, with *r*_*i*_ referring to the mean firing rate of neuron i in a population of N neurons for population (*S*_*P*_, CV_*P*_) or the mean firing rate in the i-th time window of N windows along time for a single neuron (*S*_*L*_, CV_*L*_).

### 5.6 ICE calculation for biophysical neuron and network models

For the firing rates of a biophysical neuron model, the information capacity *H* is calculated by the entropy on *P* (*x* = *r*_*i*_), the probability of the discrete distribution of the instantaneous firing rate *r*_*i*_ over time:

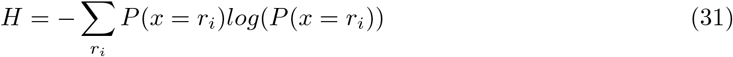

Different from the energy cost definition of a general signal in Eq.9, for a realistic biophysical neuron, we should consider its energy consumption separately for supporting processes include maintaining resting potentials, performing housekeeping functions, covering hardware wiring costs (*C*_1_), and signaling (*C*_2_ = *λC*1), then:

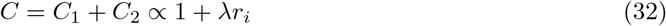

Since the total energy for signaling related demands is 75 − 78% of neuronal metabolism and nonsignaling neuronal components (including resting potential and housekeeping) occupy 22 − 25% of total neuronal metabolism [29], a rough estimate *λ* ≈ 3.5 is applied. Thus, with given distribution of simulated *r*_*i*_, we calculate ICE as:

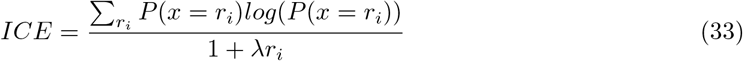

For network models, ICE is calculated for each neuron and the averaged value is used to represent the efficiency of the network so that we can compare the energy efficiency fairly for models with different amounts of neurons.

## Author Contributions

Yuguo Yu, Wei Lin, and Anna Wang Roe supervised the research, Mingyi Huang and Yuguo Yu designed the research, Mingyi Huang performed the research, Mingyi Huang deducted the proof, wrote the analysis tools and analyzed the data, Mingyi Huang and Yuguo Yu wrote the paper.

## Data Availability

Data will be made available on request.

## Supplemental Materials

Fig. S1. Monotonic relationship between the CV_*L*(*ISI*)_ and *S*_*L*(*ISI*)_.

Fig. S2. Monotonic relationship between the CV_*L*(*ISI*)_ and *S*_*L*_ for Poisson process.

Fig. S3. Monotonic relationship between the CV_*L*(*ISI*)_ and *S*_*L*_ for spike sequences with Gaussian and Uniform ISI distributions.

Fig. S4. Numerical calculation of *h*(*u*) and *g*(*u*) on *u* ∈ (0, 1). Fig. S5. Comparison of *h*(*λ*) and *e*^*λ*^ for *λ* in range (0,100).

Fig. S6. The boxplot of the structural similarity (SSIM) of the reconstructed image and ground truth and dissimilarity among basis functions *U*.

Supplemental Proof 1. Equivalence of CV_*L*_ of instantaneous firing rate and CV_*L*(*ISI*)_ for Poisson processes

Supplemental Proof 2. Positivity of *g*(*u*) on *u* ∈ (0, ∞)

Supplemental Proof 3. Monotonic relation between CV and ICE for other distributions with constant standard deviation.

