## Supplementary material for "A Unified Theory of Response Sparsity and Variability for Energy-Efficient Neural Coding": Supplemetary materials

September 26, 2024

#### 1 Supplemental Figures

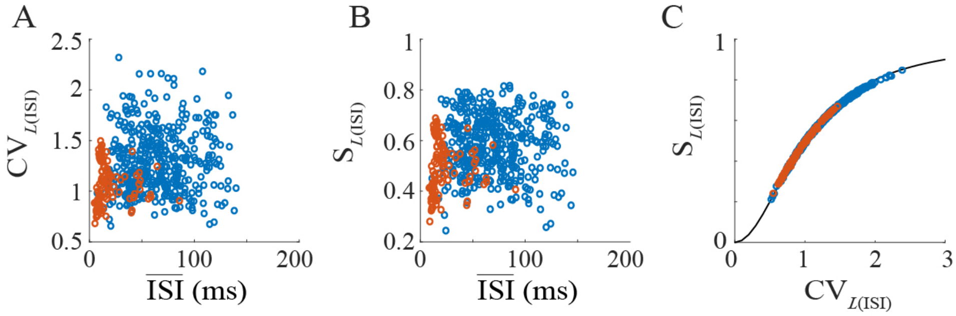


Figure S1: Monotonic relationship of between the $\mathrm{CV}_{L(ISI)}$ and $S_L(ISI)$ for experimental data (blue circle for resting state and orange for task state).

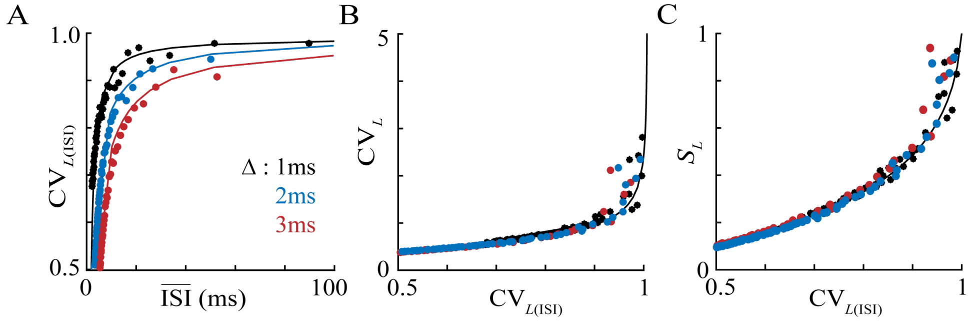


Figure S2: Monotonic relationship of between the $\mathrm{CV}_{L(ISI)}$ and $S_L$ for Poisson process with absolute refractory period. (A) Mean value of inter-spike intervals ($I\bar{S}I$) and lifetime CV calculated over inter-spike intervals ($\mathrm{CV}_{L(ISI)}$) of Poisson processes with absolute refractory periods $\Delta = 1, 2, 3ms$ (black, dark gray and light gray dots) are plotted to the analytical equation (black line). (B) $\mathrm{CV}_{L(ISI)}$ and $\mathrm{CV}_L$, which is calculated over instantaneous firing rate, are monotonically related. (C) Sparseness ($S_L$) and $\mathrm{CV}_{L(ISI)}$ of simulated processes all conform to the derived equivalence (black line).

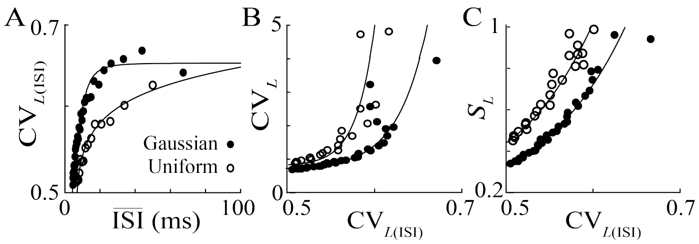


Figure S4: Monotonic relationship between the $CV_{L(ISI)}$ and $S_L$ for spike sequences with Gaussian and Uniform ISI distributions. (A) The mean value of inter-spike intervals ($\bar{ISI}$) and CV calculated over inter-spike intervals ($CV_{L(ISI)}$) for Gaussian (dots) and Uniform (circles) ISIs with a range of mean value and standard deviation controlled in the scale of corresponding Poisson sequences. (B) $CV_{L(ISI)}$ and $CV_L$, which is calculated over instantaneous firing rate, (C) Sparseness ($S_L$) and $CV_{L(ISI)}$ of simulated processes showing monotonical relations (black line).

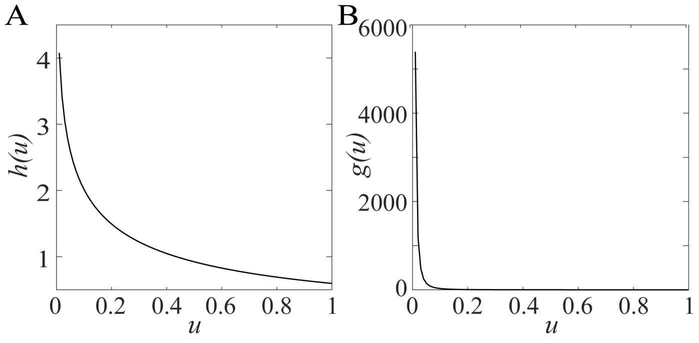


Figure S2: Numerical calculation of (A) $h(u)$ and (B) $g(u)$, suggesting the positivity on $u \in (0, 1)$.

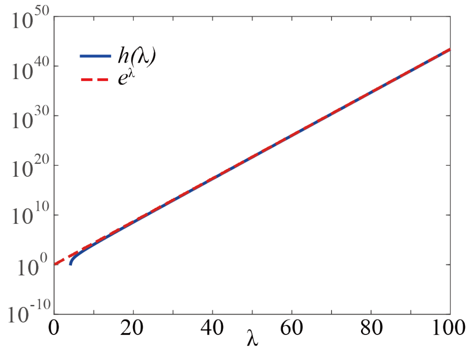


Figure S5: Comparison of $h(\lambda)$ and $e^{\lambda}$ for $\lambda$ in range (0,100) for proof in Theoretical Framework Section B. Numerical validation of $h(\lambda)$ illustrated in blue line is below $e^{\lambda}$, dashed line in red, suggesting that $h(\lambda) < e^{\lambda}$ on $\lambda \in (0, 100)$.

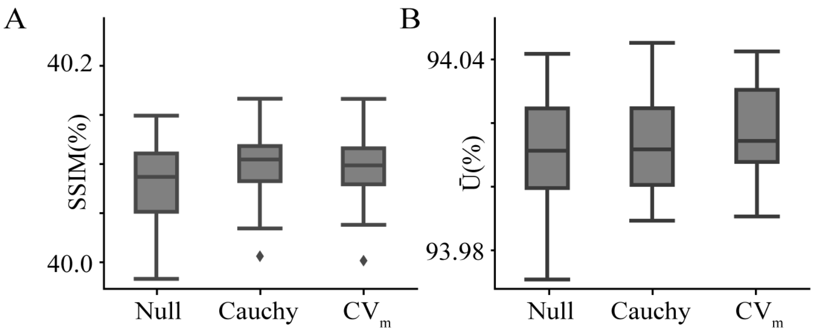


Figure S6: The boxplot compares the (A)structural similarity (SSIM) of the reconstructed image and ground truth, and (B) dissimilarity among basis functions $U$, of the null model, the sparse coding algorithm with Cauchy prior constraint and CV-maximization.

### 2 Supplemental Proof

#### 2.1 Equivalence of CV$_L$ of instantaneous firing rate and CV$_{L(ISI)}$ for Poisson processes

The interval distribution of the Poisson process with $\Delta$, the absolute refractoriness, and $R$, the mean firing rate, is given by the exponential distribution that follows:

$$P(s) = \begin{cases} 0, s \leq \Delta \\ Re^{-R(s-\Delta)}, s > \Delta \end{cases} \tag{1}$$

Then, theoretically we have $R \leq 1/\Delta$, the expectations of interval $s$ and instantaneous firing rate $r = 1/s$ can be calculated as follows:

$$\langle s \rangle = \int_{\Delta}^{\infty} Re^{-R(s-\Delta)} s ds = \frac{\Delta R + 1}{R}, \tag{2}$$

$$\langle s^2 \rangle = \int_{\Delta}^{\infty} Re^{-R(s-\Delta)} s^2 ds = \frac{\Delta^2 R^2 + 2\Delta R + 2}{R^2}, \tag{3}$$

$$\langle r \rangle = \int_{\Delta}^{\infty} \frac{R}{s} e^{-R(s-\Delta)} ds = Re^{\Delta R} \int_{\Delta R}^{\infty} \frac{e^{-t}}{t} dt, \tag{4}$$

$$\begin{aligned} \langle r^2 \rangle &= \int_{\Delta}^{\infty} \frac{R}{s^2} e^{-R(s-\Delta)} ds \\ &= R^2 e^{\Delta R} \int_{\Delta R}^{\infty} \frac{e^{-t}}{t^2} dt \\ &= R^2 e^{\Delta R} \left( \frac{e^{-\Delta R}}{\Delta R} - \int_{\Delta R}^{\infty} \frac{e^{-t}}{t} dt \right) \\ &= \frac{R}{\Delta} - R^2 e^{\Delta R} \int_{\Delta R}^{\infty} \frac{e^{-t}}{t} dt \end{aligned} \tag{5}$$

Using the above calculation results, respectively, to CV$_{L(ISI)}$ and CV$_L$ yields:

$$CV_{L(ISI)} = \sqrt{\frac{\langle s^2 \rangle}{\langle s \rangle^2} - 1} = \frac{1}{\Delta R + 1}, \tag{6}$$

$$CV_L = \sqrt{\frac{\langle r^2 \rangle}{\langle r \rangle^2} - 1}. \tag{7}$$

Here, $\mathrm{CV}_{L(ISI)}$ is denoted as $x$, with $u = \Delta R = 1/x - 1$, we consider the function $f(u) = \frac{\langle r^2 \rangle}{\langle r \rangle^2}$. Then:

$$f(u) = \frac{\frac{R}{\Delta} - R^2 e^{\Delta R} \int_{\Delta R}^{\infty} \frac{e^{-t}}{t} dt}{(R e^{\Delta R} \int_{\Delta R}^{\infty} \frac{e^{-t}}{t} dt)^2} = \frac{\frac{1}{u} - e^u \int_u^{\infty} \frac{e^{-t}}{t} dt}{(e^u \int_u^{\infty} \frac{e^{-t}}{t} dt)^2} \tag{8}$$

Denote $h(u) = e^u \int_u^{\infty} \frac{e^{-t}}{t} dt$, then $f(u) = \frac{1/u - h(u)}{h^2(u)}$, its derivative can be formatted as follows:

$$\begin{aligned} f'(u) &= \frac{1}{h^4(u)}((-\frac{1}{u^2} - h'(u))h^2(u) - 2h(u)h'(u)(\frac{1}{u} - h(u))) \\ &= -\frac{1}{h^4(u)}(\frac{h^2(u)}{u^2} + h'(u)h^2(u) + 2\frac{h(u)h'(u)}{u} - 2h^2(u)h'(u)) \\ &= -\frac{1}{h^2(u)}(\frac{1}{u^2} + h'(u)(\frac{2}{uh(u)} - 1)) \end{aligned} \tag{9}$$

As suggested by Fig. S2, $g(u) = \frac{1}{u^2} + h'(u)(\frac{2}{uh(u)} - 1) > 0$ on $u \in (0, 1)$ (for strict proof of this positivity see Supplemental proof 2.1), thus $f'(u) < 0$. Since $u = 1/x - 1$, it is sufficient to prove that $g'(x) = f'(u)u'(x) > 0$. Therefore, $\mathrm{CV}_L = \sqrt{g(x) - 1}$ is a monotonically increasing function of x, that is, $\mathrm{CV}_{L(ISI)}$.

Therefore, the equivalence between $\mathrm{CV}_L$ and $\mathrm{CV}_{L(ISI)}$ stands. Then, we further obtain the sparseness as a monotonic increasing function of $\mathrm{CV}_{L(ISI)}$, which is denoted by $x$:

$$S = 1 - \frac{(1-x)e^{\frac{2}{x}} \left( \int_{\frac{1}{x}-1}^{\infty} \frac{e^{-t}}{t} dt \right)^2}{e^2 x - e^{\frac{x+1}{x}} (1-x) \int_{\frac{1}{x}-1}^{\infty} \frac{e^{-t}}{t} dt}. \tag{10}$$

#### 2.2 Positivity of $g(u)$ on $u \in (0, \infty)$

*Proof.* Recall that

$$h(u) = e^u \int_u^{\infty} \frac{e^{-t}}{t} dt \tag{11}$$

$$h'(u) = e^u \int_u^{\infty} \frac{e^{-t}}{t} dt - \frac{1}{u} = h(u) - \frac{1}{u} \tag{12}$$

Thus we have $g(u)$ as:

$$\begin{aligned} g(u) &= \frac{1}{u^2} + h'(u)(\frac{2}{uh(u)} - 1) \\ &= \frac{1}{u^2} + (h(u) - \frac{1}{u})(\frac{2}{uh(u)} - 1) \\ &= \frac{1}{u^2} + \frac{2}{u} - h(u) - \frac{2}{u^2 h(u)} + \frac{1}{u} \\ &= \frac{1}{u^2 h(u)}(h(u) - 2 + 3uh(u) - u^2 h^2(u)) \end{aligned} \tag{13}$$

Since $h(u)$ as well as $uh(u)$ is intuitively positive (numerical calculation see Fig. S2A), showing that $\lim_{u\to\infty} uh(u) = 1$ is sufficient for proving $g(u) > 0$. First, the following equality of the first-order exponential integral over real variables stands:

$$\int_u^\infty \frac{e^{-t}}{t} dt = \int_{-\infty}^{-u} -\frac{e^v}{v} d(-v) = \int_{-\infty}^{-u} \frac{e^v}{v} dv \tag{14}$$

Then, we have

$$\begin{aligned}\lim_{u\to\infty} uh(u) &= \lim_{u\to\infty} ue^u \int_u^\infty \frac{e^{-t}}{t} dt \\ &= \lim_{u\to\infty} \int_{-\infty}^{-u} \frac{e^{-v}}{v} dv / \left(\frac{e^{-u}}{u}\right) \\ &= \lim_{u\to\infty} \frac{ue^{-u}}{e^{-u} + ue^{-u}} \\ &= \lim_{u\to\infty} \frac{u}{1+u} = 1\end{aligned} \tag{15}$$

Interestingly, recall that $u = \Delta R$, with $\Delta$ and $R$ defined as absolute refractory period and mean firing rate of a neuron, another intuitive understanding of this limit is as follows:

$$\lim_{R\to\infty} \Delta R e^{\Delta R} \int_{\Delta R}^\infty \frac{e^{-t}}{t} dt = \lim_{R\to\infty} \Delta \langle r \rangle = 1, \tag{16}$$

which suggests that when the mean firing rate R is sufficiently large, the expectation of the inter-spike interval approaches the absolute refractory period $\Delta$ and that the expectation of the instantaneous firing rate $\langle r \rangle$ approaches $\frac{1}{\Delta}$ accordingly. Hence, $\lim_{u\to\infty}(3uh(u) - u^2h^2(u) - 2) = 0$.

Therefore, as a monotonically decreasing function (see Fig. S2B), the positivity of $g(u)$ is proven by:

$$\begin{aligned}\lim_{u\to\infty} g(u) &= \frac{1}{u^2h(u)}(h(u) + 3uh(u) - u^2h^2(u) - 2) \\ &= \frac{1}{u^2} = 0.\end{aligned} \tag{17}$$

□

#### 2.3 Monotonic relation between CV and ICE for other distributions with constant standard deviation

##### 2.3.1 Uniform Distribution

Consider a set of variables that conforms to Uniform distribution on $x \in [\mu - u, \mu + u]$:

$$P(x) = \frac{1}{2u} \tag{18}$$

Then naturally, we have the standard deviation $\sigma = u/\sqrt{3}$.

Then, the entropy and cost can be calculated respectively as:

$$H = -\int_{\mu-\sqrt{3}\sigma}^{\mu+\sqrt{3}\sigma} \frac{1}{2\sqrt{3}\sigma} log(\frac{1}{2\sqrt{3}\sigma})dx = log2 + log\sqrt{3}\sigma \tag{19}$$

$$C = \mu^2 + \sigma^2 \tag{20}$$

Consider the condition where $\sigma$ is a constant, we have ICE= $f(\mu)$ as a decreasing function of $\mu \in (0, \infty)$ :

$$f(\mu) = \frac{log2 + log\sqrt{3}\sigma}{\mu^2 + \sigma^2} \tag{21}$$

Denote CV with $t$, then $t = \sigma/\mu > 0$, thus we have $f(t)$ and its derivative as:

$$f(t) = \frac{log2 + log\sqrt{3}\sigma}{\sigma^2} \times \frac{t^2}{t^2+1} f'(t) = \frac{log2 + log\sqrt{3}\sigma}{\sigma^2} \times \frac{2t}{(t^2+1)^2} > 0 \tag{22}$$

Therefore, ICE increases along with CV for variables that conform to a uniform distribution.

##### 2.3.2 Gaussian Distribution

Consider a set of variables that conforms to Uniform distribution on $x \in N(\mu, \sigma^2)$:

$$P(x) = \frac{1}{\sqrt{2\pi}\sigma} e^{-\frac{(x-\mu)^2}{2\sigma^2}} \tag{23}$$

Then, the entropy and cost can be calculated respectively as:

$$\begin{aligned} H &= -\frac{1}{\sqrt{2\pi}\sigma} \int_{-\infty}^{\infty} e^{-\frac{(x-\mu)^2}{2\sigma^2}} \log\left(\frac{e^{-\frac{(x-\mu)^2}{2\sigma^2}}}{\sqrt{2\pi}\sigma}\right) dx \\ &= \frac{1}{\sqrt{2\pi}\sigma}\left(\log\left(\sqrt{2\pi}\sigma\right) + \int_{-\infty}^{\infty} \frac{(x-\mu)^2}{2\sigma^2} e^{-\frac{(x-\mu)^2}{2\sigma^2}} dx\right) \\ &= \frac{1}{\sqrt{2\pi}\sigma}\left(\log\left(\sqrt{2\pi}\sigma\right) + \frac{1}{2}\right) \end{aligned} \tag{24}$$

$$C = \mu^2 + \sigma^2 \tag{25}$$

ICE= $f(\mu)$ as a decreasing function of $\mu \in (0, \infty)$ :

$$f(\mu) = \frac{\frac{1}{\sqrt{2\pi}\sigma}\left(\log\left(\sqrt{2\pi}\sigma\right) + \frac{1}{2}\right)}{\mu^2 + \sigma^2} \tag{26}$$

Denoting CV with $t$, then $t = \sigma/\mu > 0$, we have ICE as a function $f(t)$ along with its derivative:

$$f(t) = \frac{\left(\log\left(\sqrt{2\pi}\sigma\right) + \frac{1}{2}\right)}{\sqrt{2\pi}\sigma^3} \times \frac{t^2}{t^2+1} f'(t) = \frac{\left(\log\left(\sqrt{2\pi}\sigma\right) + \frac{1}{2}\right)}{\sqrt{2\pi}\sigma^3} \times \frac{2t}{(t^2+1)^2} > 0 \tag{27}$$

Since $\sigma$ is also a constant, $f(t)$ is a monotonically increasing function of $t$, that is, ICE is a monotonically increasing function of CV. Therefore, ICE increases along with CV for variables from a Gaussian distribution.
